# Rabix: an open-source workflow executor supporting recomputability and interoperability of workflow descriptions

**DOI:** 10.1101/074708

**Authors:** Gaurav Kaushik, Sinisa Ivkovic, Janko Simonovic, Nebojsa Tijanic, Brandi Davis-Dusenbery, Deniz Kural

## Abstract

As biomedical data has become increasingly easy to generate in large quantities, the methods used to analyze it have proliferated rapidly. Reproducible and reusable methods are required to learn from large volumes of data reliably. To address this issue, numerous groups have developed workflow specifications or execution engines, which provide a framework with which to perform a sequence of analyses. One such specification is the Common Workflow Language, an emerging standard which provides a robust and flexible framework for describing data analysis tools and workflows. In addition, reproducibility can be furthered by executors or workflow engines which interpret the specification and enable additional features, such as error logging, file organization, optim^1^izations tocomputation and job scheduling, and allow for easy computing on large volumes of data. To this end, we have developed the Rabix Executor^a^, an open-source workflow engine for the purposes of improving reproducibility through reusability and interoperability of workflow descriptions.

## 1. Introduction

Reproducible analyses require the sharing of data, methods, and computational resources.^1^ The probability of reproducing a computational analysis is increased by methods that support replicating each analysis and the capability to reuse code in multiple environments. In recent years, the practice of organizing data analysis via computational workflow engines or accompanying workflow description languages has surged in popularity as a way to support the reproducible analysis of massive genomics datasets.^2,3^ Robust and reliable workflow systems share three key properties: flexibility, portability, and reproducibility. Flexibility can be defined as the ability to gracefully handle large volumes of data with multiple formats. Adopting flexibility as a design principle for workflows ensures that multiple versions of a workflow are not required for different datasets and a single workflow or pipeline can be applied in many use cases. Together, these properties reduce the software engineering burden accompanying large-scale data analysis. Portability, or the ability to execute analyses in multiple environments, grants researchers the ability to access additional computational resources with which to analyze their data. For example, workflows highly customized for a particular infrastructure make it challenging to port analyses to other environments and thus scale or collaborate with other researchers. Well-designed workflow systems must also support reproducibility in science. In the context of workflow execution, computational reproducibility (or recomputability) can be simply defined as the ability to achieve the same results on the same data regardless of the computing environment or when the analysis is performed. Workflows and the languages that describe them must account for the complexity of the information being generated from biological samples and the variation in the computational space in which they are employed. Without flexible, portable, and reproducible workflows, the ability for massive and collaborative genomics projects to arrive at synonymous or agreeable results is limited.^4,5^

Biomedical or genomics workflows may consist of dozens of tools with hundreds of parameters to handle a variety of use cases and data types. Workflows can be made more flexible by allowing for transformations on inputs during execution or incorporating metadata, such as sample type or reference genome, into the execution. They can allow for handling many use cases, such as dynamically generating the appropriate command based on file type or size, without needing to modify the workflow description to adjust for edge cases. Such design approaches are advantageous as they alleviate the software engineering burden and thus the accompanying probability of error associated with executing extremely complex workflows on large volumes of data. In addition, as the complexity of an individual workflow increases to handle a variety of use cases or criteria, it becomes more challenging to optimally compute with it. For example, analyses may incorporate nested workflows, business logic, memoization or the ability to restart failed workflows, or require parsing of metadata -- all of which compound the challenges in optimizing workflow execution.

As a result of the increasing volume of biomedical data, analytical complexity, and the scale of collaborative initiatives focused on data analysis, reliable and reproducible analysis of biomedical data has become a significant concern. Workflow descriptions and the engines that interpret and execute them must be able to support a plethora of computational environments and ensure reproducibility and efficiency while operating across them. It is for this reason that we have developed the Rabix Executor (on GitHub as Project “Bunny”)^a^, an open-source workflow engine designed to support computational reproducibility/recomputability through the use of standard workflow descriptions, a software model that supports metadata integration, provenance over file organization, the ability to reuse workflows efficiently, and which combines an array of optimizations used separately in existing workflow execution methods.^6–12^

For the 1.0 release of the Rabix Executor (or Rabix), we’ve focused on supporting the Common Workflow Language (CWL), an open, community-driven specification for describing tools and workflows with a focus on features that support reproducibility.^2^ The Common Workflow Language is used to describe individual “processes” or “applications”, which can be either a single tool or an entire workflow. Workflows are described as a series of “steps,” each of which is a single tool or another, previously-described workflow. Each step in the workflow has a set of “ports” which represent data elements that are either inputs or outputs of the tool. A single port represents a specific data element that is required for execution of the tool or is the result of its execution. For data elements that are passed between applications, there must be an output port from the upstream application and a complementary input port on the downstream application.

CWL is designed to be extensible, so the specification may grow based on the community’s needs. However, the software model for Rabix was designed with an abstract workflow execution model to anticipate support for additional workflow languages or syntax used by other workflow engines.

## 2. Software model used by Rabix to interpret and compute workflows

The Rabix Executor allows users to execute applications described by a workflow description language. First, the workflow description is submitted to the engine. Then, the Rabix engine interprets the workflow description and translates it into discrete computational processes or “jobs.” Finally, the jobs are queued to a backend or computational infrastructure, such as a local machine, cluster, or cloud instances, for scheduling and execution. Each component of the executor (frontend, bindings, engine, queue, backend) is abstracted from each other to enable complete modularity; Developers are able to design custom frontends (e.g. command line or graphical user interface), bindings for the engine to parse different workflow languages, use the queuing protocol of their choice, and submit computational jobs to different backends. This flexible software model means that Rabix can be modified to perform data analysis on many different infrastructures as desired by the user or developer and achieve identical results or incorporate tools described by different languages or syntaxes into a single workflow.

## 3. Abstract representation of data analysis workflows in Rabix

Computational workflows are frequently understood as a directed acyclic graph (DAG)^3,13,14^, a kind of finite graph which contains no cycles and which must be traversed in a specific direction. In this representation, each node is either an individual executable command, a “nested” workflow, or a set of commands that can be executed in parallel. The edges in the DAG represent execution variables (data elements such as files or parameters) which pass from upstream nodes to downstream ones.

**Figure 1.**
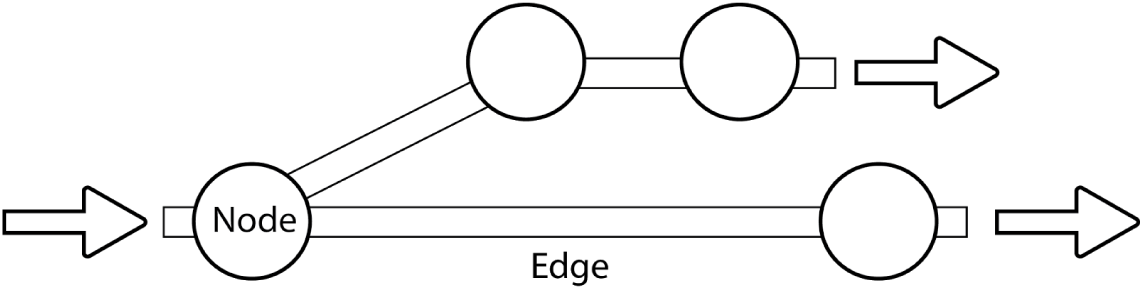
Illustration of a directed acyclic graph (DAG). The DAG may be traversed from left-to-right, moving from node-to-node along the edges that connect them.

Workflows can be described as machine-readable serialized data objects in either a general-purpose programming language (GPL), domain-specific language (DSL), or serialized object models for workflow description.^2,9,15^ For example, an object model-based approach may describe the steps in a workflow in JSON format with a custom syntax. This workflow description can then be parsed by an engine or executor to create the DAG representation of the workflow. The executor may then translate the directions for workflow execution to actionable jobs in which data is analyzed on a computational infrastructure, such as a cloud computing instance, a high-performance computing cluster, or a personal computer.

A primary design constraint of the Rabix executor is to abstract components of a workflow to a data model that is comprehensive enough to allow for mapping the syntax of different workflow systems, whether they are DSLs or serialized data objects. In this way, tools and workflows from different systems can be used together in a single workflow.

### 3.1. General structure of a workflow execution

There are three general steps in preparing a workflow for execution: interpretation of a machine-readable workflow description, generation of the workflow DAG, and finally decomposition into individual jobs that can be scheduled for execution. At the beginning of execution, a workflow engine or interpreter is provided with the workflow description and the required inputs for execution of the workflow, such as parameters and file paths (Fig. 2a). The workflow description object is then parsed and a DAG is created (Fig. 2b), which contains the initial set of nodes and edges required for computation.

**Figure 2.**
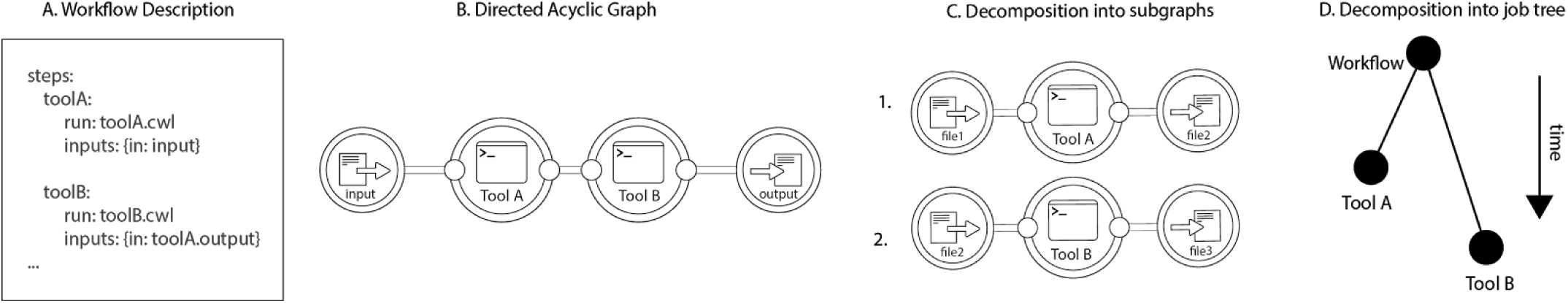
The process of parsing a workflow description. **A.** The machine-readable document is interpreted, from which **B.** a DAG is produced. From the DAG, **C.** subgraphs representing computational jobs that can be sent to backends for scheduling/execution and **D.** a job tree is resolved, which identifies “parent” and “leaf” nodes. Each leaf represents an individual job.

In addition to representing the steps in the workflow as a DAG (Fig. 2c), certain workflow ontologies model computational jobs as a composite (tree) pattern in which there are “parent nodes” (workflows), which can contain multiple executables or “leaf nodes” or other "parent" nodes (Fig. 2d).^16–20^ The Rabix engine extends this model by generalizing “parent” nodes to include groups of jobs, such as when parallelization is possible at that node. It is important to note that the “parent-child” terminology is also applied to relations between individual workflow nodes by the Toil project, an executor which can also interpret Common Workflow Language.^10^ However, Rabix uses these terms to refer to computational "jobs" and "subjobs", e.g. a “nested” workflow node is a child of a workflow and can be decomposed into an array of “subjobs”. The engine handles the "execution" or parsing of these parent jobs, while leaves are queued for scheduling and execution on a backend. This model allows for more efficient resolution of DAG features such as nodes in which steps can be parallelized or are nested. It also maintains a one-to-one mapping between the internal DAG representation and the workflow description supplied by the author.

## 4. Optimization of CWL workflows via DAG transformations

The Rabix Executor began its development by examining how to interpret Common Workflow Language and interoperate on different versions or earlier drafts, in a way that is extensible to future versions and other workflow syntaxes. Rabix currently supports tools and workflows described in CWL Draft 2, Draft 3, and version 1.0, either individually or in combination.

When a CWL workflow is represented as a DAG, applications become nodes and edges indicate the flow of data elements between ports of linked tools. In the case of a simple workflow, there are no possible transformations of the DAG; each node represents a single command line execution and all data elements are simply passed from tool-to-tool as-is (Fig. 3).

**Figure 3.**
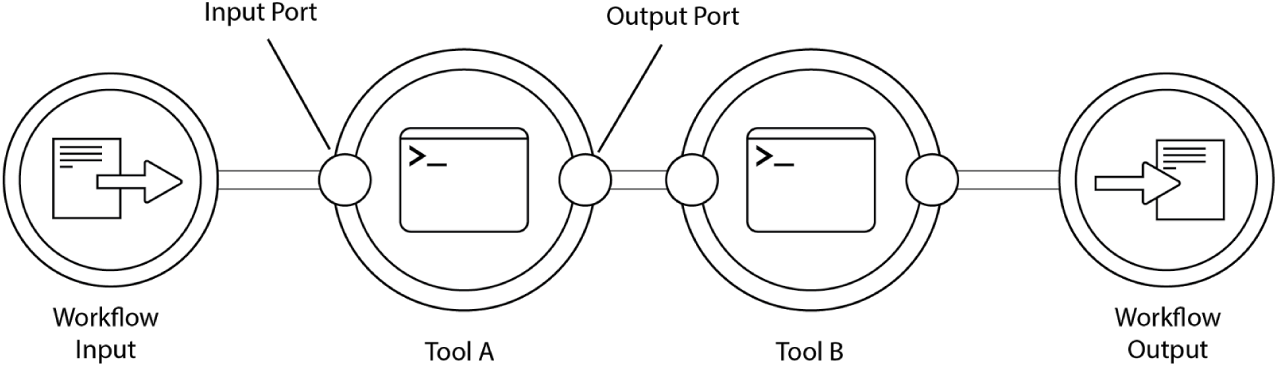
A DAG created from a workflow described by the Common Workflow Language which contains two tools (**A**, **B**). Tools have input and output ports, which define discrete data elements that are passed downstream along the edges of the DAG.

Additionally, CWL workflows can be designed such that data elements and the execution itself can be transformed during runtime. Developers are given several options for describing workflows which can enhance their utility and flexibility in handling biomedical data analysis:

1. The ability to generate “dynamic expressions” or transformations on data elements, inputs, outputs, and other command line arguments.
2. The ability to perform “scatter/gather I/O (input/output)”, also known as vectored I/O, in which execution of the input data can be parallelized based on specific criteria. A common genomics use case for this is performing an analysis per chromosome, in which the set of chromosomes is delivered to a node as an array (e.g. [1, 2, 3, X]).
3. The ability to nest workflows within workflows, which allows for rapid composition of complex workflows and the ability to quickly reuse existing code.

### 4.1 Rabix uses a custom data model and port-level inspection for workflow execution

Though CWL provides a specification for how to describe the execution of tools and workflows, the exact way in which these features are implemented is left entirely to the execution engine that is interpreting it. Therefore, the Rabix engine has been designed to handle CWL descriptions with two optimizations:

1. Reacting to "port ready" events rather than "job done" events. “Port ready” is a state triggered by the evaluation of data elements produced by a port, whereas “job done” refers to *all* ports of a node being evaluated. In this approach, possible downstream executions are triggered if the edges leading to it are resolved. This allows further dynamic transformations of the DAG to optimize for when all prerequisites for downstream jobs are ready.

2. Reacting to "port ready" events from dynamically created subjobs and rewiring them to their final destinations, possibly creating and running subjobs before their parent fully evaluated (referred to as “look-ahead” method).

These functionalities enable the Rabix engine to create additional edges and nodes as needed, in order to decompose the workflow DAG as early as possible, allowing downstream jobs to be scheduled as soon as actual prerequisites are met.

The workflow DAG is stored in three tables, Variables, Jobs, and Links, which are accessed when a port value is updated. The Variables table contains the ports and their explicit values. The Jobs table stores each node of the workflow and a counter for the inputs and outputs that have been evaluated at that node. The Links table stores the edges in the DAG that is traversed.

As compared to other CWL execution models^2,10^, computational events are triggered by “port” events instead of “job” events. In other words, when a port is evaluated, this triggers the executor to scan or update these tables in the following order: Variables, Jobs, Links. Any node for which all input ports are now evaluated is then executed.

Suppose for example, Rabix is executing the workflow in Figure 4. The engine will first parse the workflow description as a workflow DAG with two variables (W.I, W.O; Fig. 4a), which are yet to be evaluated. Additionally, there are two ports (#In, #Out), an input and an output. Next, the engine inspects the contents of the workflow (Fig. 4b) and is able to see the following steps: Tool A, Tool B, each of their ports, and the link between each step within scope.

**Figure 4.**
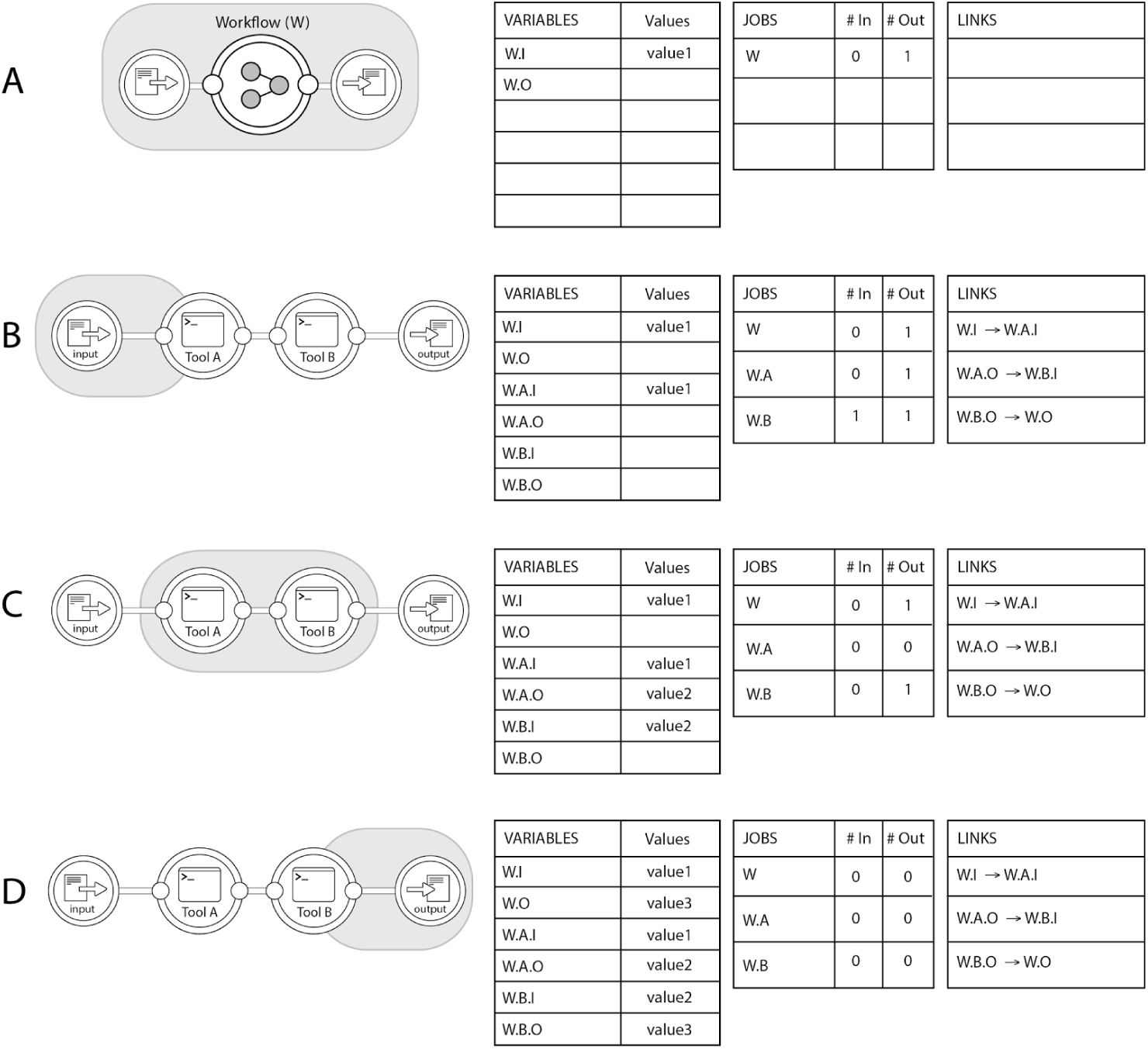
The algorithm as it is traversed. **A.** The engine interprets the top-level of the workflow description and **B.** inspects the contents of the workflow node and determines the DAG structure and links between each step (edges). The currying of value1 from the workflow input to the input of Tool A triggers an input event, where a job (analysis of Tool A with its inputs) is sent to a backend node. **C.** The execution continues and the engine traverses the DAG. **D.** The workflow is completed when the output of the final tool (W.B.O., value3) is curried to the overall workflow output (W.O). The port counters allow the engine to track when nodes are ready to be executed even if upstream jobs are only partially completed.

After this, any known values are curried downstream through their links. The input for the workflow (W.I) is curried to Tool A through the link that has been identified between the two (W.I → W.I.A). The input job counter (#In) for Tool A is decremented to 0, thereby triggering an input event where a job (execution of Tool A with value1) is distributed to a backend for computation. The engine now waits for an event in which the output of Tool A (W.A.O) is reported as value.

Once the output for the job is evaluated and reported to the engine (value2), an output event is triggered. The output port for W.A is decremented to 0, the link from W.A.O to W.B.I is traversed, and W.B.I is evaluated as value2. This reduces the #In counter for W.B to 0 in the Jobs table and triggers a job, the execution of Tool B with its input (Fig. 4c). The execution finally concludes until the input port counter for W reaches 0 and W.O is evaluated (Fig. 4d).

In the case where the engine is traversing a portion of the workflow that maps to a parent node beneath the root parent node, each output update event will generate an additional output update event. This strategy allows the engine to “look-ahead” towards future executions and apply optimizations to dynamic portions of the DAG, as outlined in the following sections.

### 4.2. DAG transformations: parallelization with scatter/gather

By evaluating workflows through this port-counter and trigger system, Rabix is capable of rewiring parallelizable nodes in the DAG when upstream jobs are only partially completed. Suppose we have a workflow where a data file and an array are inputs for a single tool, which then produces an output file (Fig. 5a). In this case, the tool is capable of being scattered over an array of variables (e.g. [1, 2, 3]). Normally, these executions will be performed sequentially on a single core, or on multiple threads if the tool allows it. However, on a workflow level, additional parallelization can be enabled by scattering the data over three separate executions of the tool based on the values in the array (Fig. 5b), thus allowing the jobs to be distributed to separate computational instances as needed.

**Figure 5.**
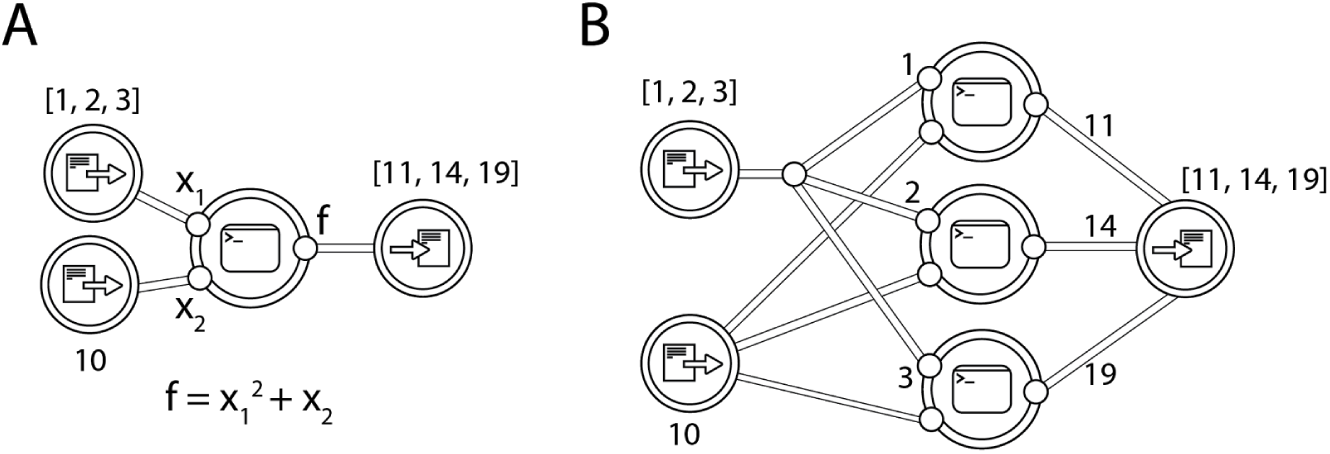
Graph transformations when performing parallelization. In this workflow, a function is performed on two inputs, an *int* and an *array of ints*. **B.** The flattened DAG created by the engine. Each value of the array is scattered as a single process to reduce computation time.

The advantages of the transformation approach is further demonstrated by another use case, in which there are two sequential, parallelizable jobs (Fig. 6a). Rabix employs a “look-ahead” strategy (Fig. 6b) which can mark downstream jobs as ready even though not all sub-jobs (leaves) are done from the upstream parent job.

**Figure 6.**
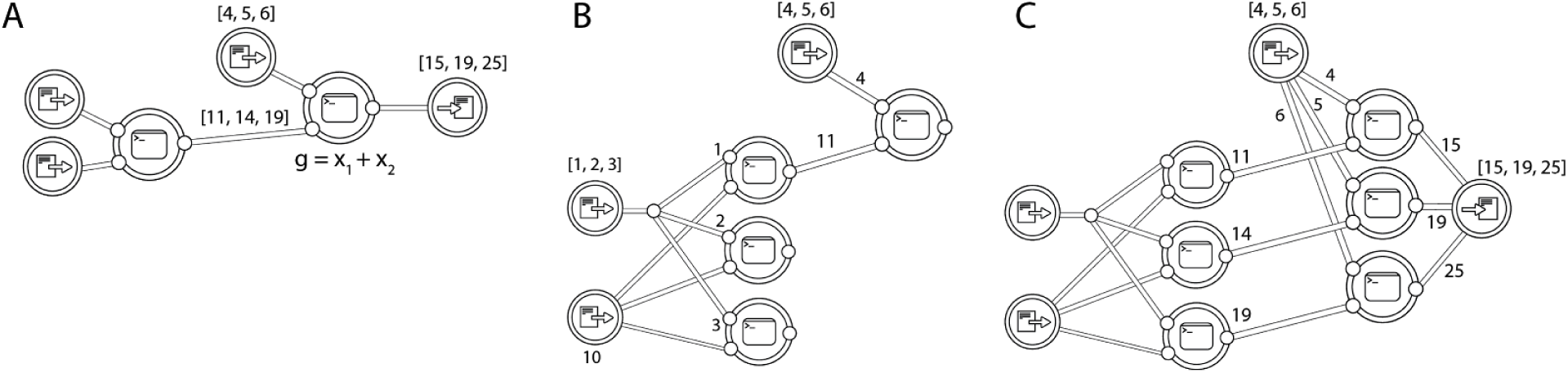
Graph transformations for sequential scattered nodes. **A.** The workflow from Fig. 5 with an additional downstream function with an input that can be scattered. **B.** During execution, the engine is able to look ahead to the next stage in the workflow. If any input is available (e.g. value of 11 returned by a tool), downstream processes which can proceed are started. **C.** The completed workflow.

Each node in the DAG does not need to be scheduled independently. Instead, (sub)jobs that work with same data can be explicitly dispatched to the same backend. (Fig. 7). For example, in the case of executions scattered across chromosome number, jobs processing the same chromosome can be distributed to the same node to optimize cost.

**Figure 7.**
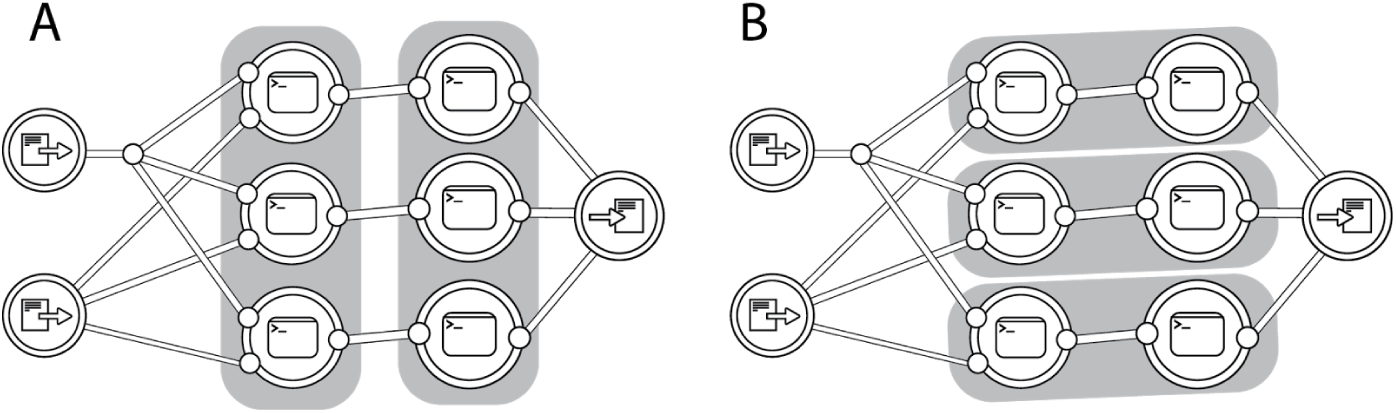
Jobs can be grouped (grey background) for execution on a backend node from criteria set by the workflow or tool author.

Figure 7 demonstrates two possible job group assignments. In the case of Figure 7a, the first tool can be executed simultaneously for each chunk of the data on a single backend node. Once any single job in the first group is finished, the second group of jobs can begin execution on a second node. In the case of Figure 7b, each chunk of data is parallelized across three nodes and the final output is gathered at the end.

The engine is also able to send information to the backends about upcoming jobs, which allows a backend scheduler to pre-allocate resources for them. When executing CWL workflows, both of these optimizations are enabled through the "hints" feature.

Whether these optimizations can be used are sometimes dependent on how the workflow is constructed. For example, a workflow author can make use of optimizations in Fig. 6 by grouping nodes that can be scattered into a nested workflow. This optimization can be especially useful when combined with nested workflow optimizations described in the next section, and allows for reusability of previously made workflows, as encouraged by CWL.

### 4.3. Graph transformations: nested workflows

CWL developers have the ability to reuse existing code and import previously-described workflows into other workflows. This feature means that it is possible to reuse code for additional workflows in lieu of refactoring and potentially introducing errors that break reproducibility. However, the ability to nest workflows presents a challenge to interpretation and optimization by the engine. If no DAG transformations are applied and nested workflows are only executed recursively, this can lead to unnecessarily prolonged execution time and cost.

Suppose a developer has described a workflow that takes two inputs and produces two outputs from two tools (Fig. 8a). In this workflow, one of the outputs is created by the upstream tool and one from the downstream tool. Later, the developer wishes to reuse this workflow description in another workflow, where the output of the upstream tool is passed to another tool for further analysis (Fig. 8b). As with sequentially scattered tools, the engine is capable of passing values from the nested workflow, once they’re produced, to steps downstream using the “look-ahead” strategy. Commonly, the tool outside the nested workflow is blocked from execution until all outputs from the nested workflow are produced, leading to increased computation time and cost.

**Figure 8.**
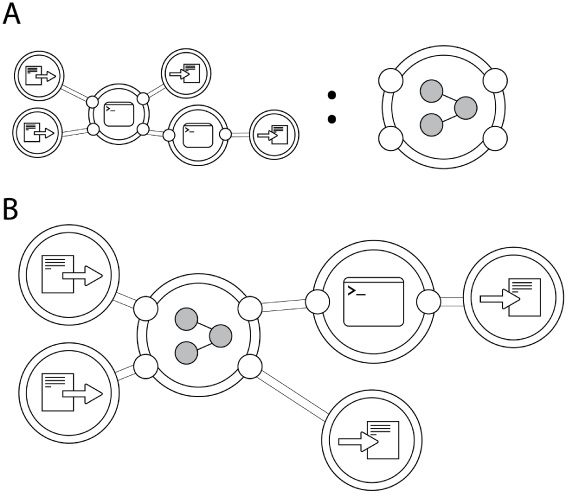
Graph transformations of nested workflows to optimize total execution time. **A.** Workflow consisting of two tools. **B.** Workflow in Fig. 8a. extended with third tool. The engine allows the downstream tool to start executing once the necessary inputs are ready, even if the upstream workflow has yet to produce all of its outputs. No code refactoring from the workflow in 8a is required.

### 4.4. Benefits to Logging, Orchestration, and Computation

The model used by the Rabix engine allows for improved optimization of data analysis at the workflow level. Further, it provides the ability to implement additional optimizations or features to enhance orchestration of jobs and computation, regardless of whether such features are supported by a workflow description language or specification.

Rabix keeps track of all jobs executed from the workflow and caches results. In addition, each parameter of a job is recorded and automatically logged for the researcher. These include the explicit command line arguments used, the files/paths, attributes of the data, metadata attributes, and any logs associated with the execution. In addition, a snapshot of the application is stored, along with the explicit values used in the execution. All of this is done at the job level, allowing for granular replication of subsets of a workflow. If the workflow contains a job that has previously been executed and the outputs are still available, the engine can reuse them even if the job was part of a different workflow run. Importantly, even if cached results are not available, the engine will look ahead in the DAG and may encounter cached downstream jobs which do have these files available, and so can resume failed or modified workflow jobs. This makes the caching mechanism comparable to declarative workflow description such as GNU Make.^8^

Additional business logic outside of a workflow specification can also be implemented. For example, CWL does not yet allow for conditional workflows, in which the entirety of DAG is not necessarily traversed but only paths based on checkpoints during the execution. Additionally, though a DAG is acyclic, Rabix could in principle enable loops for a tool or workflow which use iterative operations.

### 4.5. Caveats to Graph Transformations and Possible Solutions

An important caveat for these optimizations are external transformations in which the structure of data elements is modified before execution and thus cannot be anticipated by the engine. For example, CWL and other workflow description languages allow for modifications of input types before tool execution. In certain cases, such as for a tool which can be scattered, the data type may change or the length of the array that is being scattered cannot be known ahead of time. If the engine is unable to anticipate the length of an array that must be scattered upon execution, it is impossible for it to re-wire the DAG before evaluation. However, such hurdles can be overcome by allowing users to either define a mapping for individual array items or declaratively specifying the method of combining multiple ports before scattering (cross-product or dot-product). In these cases, the engine can still maintain its look-ahead optimizations.

### 4.6. Furthering reproducibility by extending CWL to execution descriptions

Workflows described using the Common Workflow Language require two objects for execution: the description of an application and an input object specifying the explicit values of the required inputs. Recording a task that has been previously executed is not, however, within the scope of CWL. However, an analyst may want to reinspect a prior analysis, reuse a workflow with a specific set of parameters on new data, or reanalyze the same data with a different workflow version. It is for these reasons that we have enabled an additional layer of task description and annotation within Rabix, alleviating the burden of logging the workflow execution.

Following the execution of a workflow, additional outputs and logs are produced by Rabix as a matter of course. The explicit command line execution, an object describing the output of the execution, and a description of the workflow execution are all recorded. From these objects, it is directly possible to reproduce a prior analysis or reanalyze additional data with the exact same parameters as previous. Rabix allows for replication of a previous execution or reproduction an exact workflow on new data with a single command line call. In this way, it is possible for an analyst to not only publish a workflow but also the explicit tasks as plain text files. These functionalities can be extended with new modules or plugins to enable a variety of use cases centered on reproducibility.

## 5. Rabix in the context of existing workflow models and engines

The primary design guideline for Rabix was to support Common Workflow Language in a way which will allow for supporting additional workflow languages, whether they are domain-specific languages or object-based. Further, “tools” or workflows described in different syntaxes should be interoperable such that a single workflow may be comprised of tools and workflows from a variety of syntaxes. In effect, certain optimizations described in Rabix above have been implemented in other systems, but not yet in a single executor capable of supporting emerging standards.

Most of the focus in this paper was on port-level inspection, an abstract data model for tools and workflows, and how they can enable additional optimizations when used in conjunction. However, certain features described here are also used by existing workflow systems,^6,7,10–12^ most notably the support for multiple infrastructures. Additionally, there are certain features not yet implemented in Rabix but which are seen in other systems, such as conditional steps in a workflow, as seen in Toil. Though the Rabix model allows for conditional operations (e.g. for, if, while), we chose to focus on features supporting reusability and interoperability and computational optimizations for this manuscript.

## 6. Conclusions

The Rabix Executor is an open-source project designed to enable scalable and reproducible analysis of portable workflows, which is available on GitHub (http://github.com/rabix/bunny). Computational reproducibility, the ability to replicate a prior analysis or reuse prior workflows on new data, is required for accurately judging scientific claims or enabling large-scale data analysis initiatives in which synonymous results can be compared.^4,5,21^ The Rabix engine additionally aims to optimize workflow executions by intelligently interpreting and handling complex workflows. This is achieved through a composite model in which workflows can be more fully decomposed. Finally, additional logic can be applied to optimize for user-defined variables, such as cost or execution time, regardless of the workflow description language being interpreted.

## Footnotes

a The Rabix Executor is available on GitHub: http://github.com/rabix/bunny

1 This project has been <u>funded</u> in whole or in part with Federal funds from the National Cancer Institute, National Institutes of Health, Department of Health and Human Services, under Contract No. HHSN261201400008C.

